# Segmented MS/MS acquisition of a1 ion-based strategy for in-depth proteome quantitation

**DOI:** 10.1101/2022.07.27.501662

**Authors:** Zhiting Wang, Chao Liu, Songduo Wang, Xinhang Hou, Pengyun Gong, Xiao Li, Zhen Liang, Jianhui Liu, Lihua Zhang, Yukui Zhang

**Author notes:** These authors contributed equally to this work (Zhiting Wang, Chao Liu). Correspondence should be addressed to Lihua Zhang or Jianhui Liu.

## Abstract

In-depth proteome quantitation is of great significance for understanding protein functions, advancing biological, medical, environmental and metabolic engineering research. Herein, benefiting from the high formation efficiencies and intensities of dimethyl-labeled a1 ions for accurate quantitation, we developed an in-depth a1 ion-based proteome quantitation method, named deep-APQ, by a sequential MS/MS acquisition of the high mass range for identification and the low mass range for a1 ion intensity extraction to increase quantitative protein number and sequence coverage. By the analysis of HeLa protein digests, our developed method showed deeper quantitative coverage than our previously reported a1 ion-based quantitation method without mass range segmentation and lower missing values than widely-used label-free quantitation method. It also exhibited excellent accuracy and precision within a 20-fold dynamic range. We further integrated a workflow combining the deep-APQ method with highly efficient sample preparation, high-pH and low-pH reversed-phase separation and high-field asymmetric waveform ion mobility spectrometry (FAIMS) to study *E. coli* proteome responses under the nutritional conditions of glucose and acetate. A total of 3447 proteins were quantified, representing 82% of protein-coding genes, with the average sequence coverage up to 40%, demonstrating the high coverage of quantitation results. We found that most of the quantified proteins related to chemotaxis were differentially expressed, including the low-abundance proteins such as trg, fliL, and cheA, indicating that chemotaxis would play an important role for *E. coli* cell to survive from acetate toxicity. The above results demonstrated that the deep-APQ method is of great promising to achieve the deep-coverage proteome quantitation with high confidence.

**GRAPHICAL ABSTRACT:** 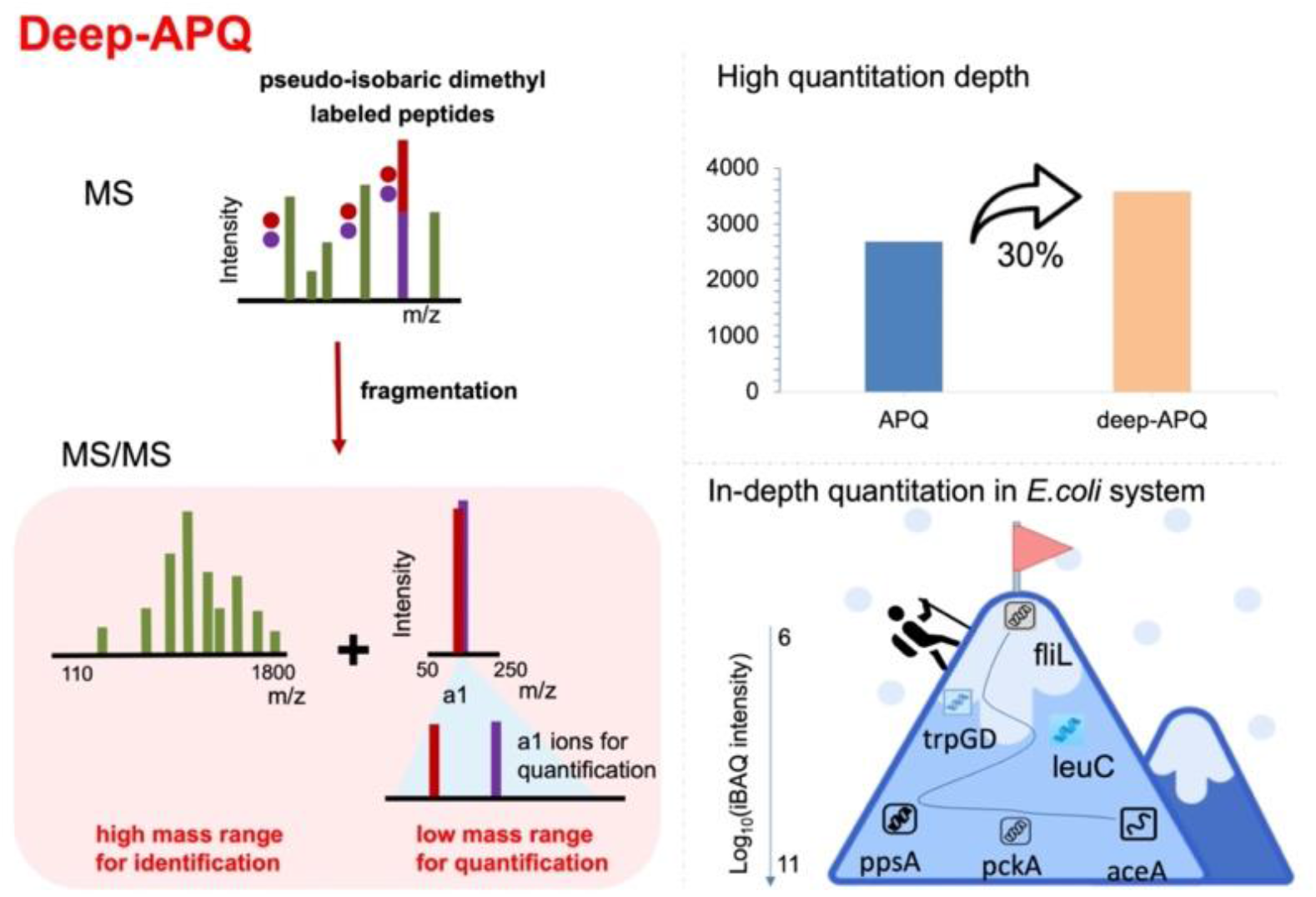

## 1. Introduction

Global proteomic quantitation, in which protein changes in response to environmental perturbations, spatial distributions and developmental states are analysed^1^, has been an effective strategy to reveal the laws of protein function and cell life activities for biology, medicine and metabolic engineering researches. Among all the quantified proteins, many proteins with low expression abundances have great biological significance, such as cell signaling, transport, cell adhesion, catalysis, and energy production. Unfortunately, the low abundance proteins are easily missed for quantitation due to the wide dynamic range of protein expressions in the proteomic samples^2^. Therefore, the in-depth quantification method is urgently needed to comprehensively analyze dynamic proteomes.

Label-free quantitation (LFQ) and labeling-based strategies are commonly used for MS-based proteome quantitation. LFQ methods analyze each sample in a single run and extract the intensities of precursor ions^3^ or fragment ions^4^ among MS data for comparison. They have been wildly used in clinical proteome studies due to the unlimitation of sample numbers and the diversity of data-acquisition schemes and algorithms. However, the low-aboundance proteins are easily missed during MS acquisition or signal extraction^5, 6^, limiting the quantitative depth. Labeling-based strategies allow quantitation of multiple samples in individual LC-MS runs by isotopic labeling and show good compatibility with multidimensional separation to greatly increase quantitative depth. In particular, reporter ion-based methods, such as tandem mass tags (TMT)^7^ and relative and absolute quantification (iTRAQ)^8^, have become popular to dynamically monitor biological processes due to its high throughput and precision. However, since the co-isolation of interference peptides and target peptide, the intensities of reporter ions are distorted, leading to ratio compression^9^.

Due to the specific masses of fragment ions that can differ the co-isolated peptides^10^, the fragment-ion-based strategies were developed to improve quantitation accuracy, precision and dynamic range^11^. Most of the fragment-ion-based strategies use b and y ions as quantitation ions for intensity extraction^10, 12, 13^. In our previous work, we found a1 ions, the neutral loss of CO molecule from b1 ions, showed enhanced intensities after dimethyl labeling and its intensities were often higher than any other fragment ions in the tandem mass spectra. Their high signal-to-noise ratios make a1 ions an ideal substitute of b, y ions for improved quantitation accuracy^10, 14^. However, the low first mass of tandem mass spectra for a1 ion acquisition and the high abundances of a1 ions lead to the loss of identification rate, especially for long peptides, reducing the quantitative numbers for comparative proteomic studies.

Herein, we presented an in-depth proteome quantitation method named deep-APQ based on segmented tandem mass spectra acquisition, including high mass range for identification and the low mass range for a1 ion intensity extraction. Higher coverage could be achieved compared to traditional APQ method, without losing accuracy and precision. Furthermore, we integrated a workflow including efficient proteome extraction, 3D separation and deep-APQ analysis to achieve in-depth proteome quantitation of *E. coli* samples from glucose and acetate carbon source growth media. All the results demonstrated that our developed approach would provide an in-depth and accurate way for understanding various physiological mechanism and providing theoretical bases for biology.

## 2. Experimental section

### 2.1. Cell culture

N^+^C^+^ minimal medium^15^ (1 g K_2_SO_4_, 17.7 g K_2_HPO_4_.3H_2_O, 4.7 g KH_2_PO_4_, 0.1 g MgSO_4_.7H_2_O and 2.5 g NaCl per litre) were used as basic growth medium, supplemented with 20 mM NH_4_Cl as nitrogen source, and 20 mM glucose or 60 mM sodium acetate as carbon source, which concentrate were depended on the number of carbon in the molecule^16^.

Growth experiment of *E. coli* cells ATCC 29425 (ATCC, Manassas, USA) for biological applications were carried out in two steps: seed culture and experimental culture, the two steps all performed in 500 mL conical flask. For seed culture, cells stored in −80°C were picked and transferred to fresh LB in an air incubator at 37°C and 135 r.p.m grown overnight. Then the cultures were centrifuged at 900 r.p.m for 5 min, the growth medium was discarded and the precipitate was re-suspended with the glucose or acetate medium to an OD_600nm_ of about 0.1 measured using an LED light source (592 nm) and a photodetector controlled by a microcontroller. For experimental culture, re-suspended cells were cultured at 37°C and 135 r. p. m, trace samples were quickly taken out nearly every 2 hours to detect OD_600nm_ to draw the growth curve (Supplementary Figure 1) until *E*. coli cells reached stationary phase.

*E. coli* cells in the middle stage of exponential growth phase, which OD_600nm_ was of about 0.85 in glucose growth medium and OD_600nm_ was of about 0.8 in acetate growth medium, were harvested by washing thrice with cold 1×PBS buffer. Three biological replicates for two kinds of samples, in total of six samples were obtained for subsequent experiment.

HeLa cells (ATCC, Manassas, USA) for quantitative method establishment and evaluation were cultured in DMEM (GIBCO, Grand Island, USA) supplemented with 10% FBS, 100 U ml^-1^ of penicillin G and 100 mg mL^-1^ of streptomycin. Cells were incubated at 37°C in an aerobic environment with 5% CO _2_ and split every 3 days following a standard protocol. Cultured HeLa cells were centrifuged at 1200 rpm for 5 min to remove culture media, and collected after washed with cold 1×PBS buffer for 3 times.

### 2.2. Sample preparation

Six samples of *E. coli* cells (5 × 10^6^) and HeLa samples (5 × 10^6^) were respectively lysed and extracted with 200 μL of C_12_Im-Cl containing 1% (v/v) cocktail, sonicated 3min with pulse set at 10 and output at 130 W on ice with a sonifier (Scientz, Ningbo, China). Cell debris was precipitated by centrifuging for 30 min at 16 000 g at 4°C, and the protein concentrations were determined by BCA protein assay (Beyotime, Nantong, China). After reduced with 100 mM DTT (Millipore, Bedford, USA) at 95°C for 5 min, the proteins were transferred to 10 kDa filter devices and washed with 50 mM phosphate buffer (PB, pH 8.0) by centrifugation at 16 000 g. Subsequently, the reduced proteins were alkylated with 150 mM Iodoacetamide (Millipore, Bedford, USA) in darkness at room temperature for 30 min. The concentrates were washed thrice with 50 mM PB by centrifugation at 16 000 g and digested with sequencing grade Trypsin-Gold (Promega, Madison, USA) at the ratio of 1:30 (enzyme/protein, m/m) overnight at 37°C. The digests were obtained by centrifugation, and the filter device were washed with 50 mM phosphate buffer. The peptide concentration was determined by nanodrop (Thermo, Rockford, USA).

### 2.3. Dimethyl labeling

For comparision of deep-APQ with APQ and LFQ method, the HeLa digests were labeled by light (4% ^13^CH_2_O and 0.6 M NaBD_3_CN, 32L) or heavy (4% CD_2_O and 0.6 M NaBH_3_CN, 32H) dimethyl labeling reagents and incubated for 1h at room temperature. Then they were mixed at the ratios of 1:1.

To evaluate the accuracy and dynamic range of deep-APQ, HeLa digests were labeled by 32L and 32H respectively and mixed at the ratios of 1:1, 1:2, 1:5, 1:10 and 1:20. All the above samples were desalted by homemade C18 column and stored at −80°C before usage.

For voltage optimization of FAIMS and analysis of *E. coli* samples in two carbon resourses, the digests of *E. coli* which cultured in glucose growth medium were labeled with 32L dimethyl labeling reagents, and the digests of *E. coli* cultured in acetate growth medium were labeled with 32H. The two labeled *E. coli* digests were mixed at the ratios of 1:1. All three mixed biological replicates were prepared for next-step separation before LC-MS analysis.

### 2.4. High-pH (hpH) reverse phase fractionation

200 μg peptides of each biological replicates samples were fractionated using a liquid chromatography (Agilent, Tokyo, Japan) with a homemade reverse phase column (C18, 5 μm, 100 Å, 2.1× 150 mm i.d., Durashell, Tianjin, China) under mobile phase at pH=10 (solvent A: 100% water, ammonia was added until pH reached 10.0, B: 80% ACN, same volume of ammonia was added to solvent B as A). Peptide samples were loaded to the column and separated using the following gradient: 5−40% B (0−40 min), 40−55% B (40−45 min), 55−90% B (45−48 min), 90% B (48−54 min), and 5% B (55−60 min), and were collected every 1 min. The resulting 60 fractions were consolidated into 20 samples with equal intervals and lyophilized, and stored at −80°C before usage.

### 2.5. nLC−(FAIMS)−MS/MS Analyses

All samples above were resuspended with 0.1% formic acid (FA). NanoRPLC experiments were performed on an EASY-nLC 1200 system (Thermo Fisher Scientific, CA, USA). Peptides were separated on a homemade C18 column (5 μm, 100 Å, 150 mm×2.1 mm i.d., Durashell, Tianjin, China) with mobile phases (solvent A: 100% H_2_O with 0.1% FA; solvent B: 20% H_2_O and 80% ACN with 0.1% FA). The separation gradient for comparison of deep-APQ and APQ method was achieved by applying 8−25% B for 130 min, and 25−50% B for 40 min. The separation gradient for comparison of deep-APQ and LFQ method, voltage optimization of FAIMS, quantitative evaluation and proteomic analysis in *E. coli* system was performed by applying 8−25% B for 35 min and 25−50% B for 25 min.

Lumos mass spectrometer (Thermo, CA, USA) was operated in positive ionization mode (a spray voltage of 2200V without and with FAIMS, respectively) and DDA mode with a fixed 3 s cycle time. For comparison of deep-APQ, APQ and LFQ method and quantitative evaluation, no FAIMS was applied to. For analysis of *E. coli* samples in two carbon resourses, 40V, 60V, and 80V were used.

For all experiments, the MS1 spectra were acquired over a mass-to-charge (m/z) range of 350−1500 m/z at a resolution of 60 000 (at m/z 200) in the Orbitrap using a maximum injection time (maxIT) of 50 ms and a normalized AGC target value of 1 × 10^6^. The top 20 most intense ions were selected with the isolation window of 1.6 m/z for MS/MS events. The charge state filter of precursors was set to 2−6, and peptiedes were fragmented by higher-energy collision dissociation (HCD) with the normalized collision energy of 30%.

For MS2, the parameter settings of APQ, deep-APQ, and LFQ methods are described below. In APQ method, a mass-to-charge (m/z) range set from 50 m/z at a resolution of 30 000 (at m/z 200) with a maxIT of 80 ms and an AGC target of 1 × 10^6^. In OTIT mode of deep-APQ, fragment ions were firstly detected in the Iontrap analyzer with the mass range set from 110, within a maxIT of 35 ms and an AGC target of 1 × 10^5^. Meanwhile, fragment ions were detected in the Orbitrap analyzer at the resolution of 30 000 (at m/z 200) with the mass range from 50 m/z to 250 m/z, and a maxIT of 80 ms and an AGC target of 1 × 10^6^. In OTOT mode of deep-APQ, one of the scan event’s parameters were kept the same as the Orbitrap analyzer above, another scan range was set from 110 within a maxIT of 50 ms and an AGC target of 5 × 10^4^. In LFQ method, fragment ions were detected in the Orbitrap analyzer with the mass range set from 110 at a resolution of 15 000 (at m/z 200), within a maxIT of 80 ms and an AGC target of 1 × 10^6^.

### 2.6. MS Data Analysis

Acquired raw files were proposed by Protein Discoverer (version 2.2) using the integrated Mascot Search engine and searched against UniProt human database (Proteome ID: UP000005640; Downloaded in 08/10/20 containing 38007 reviewed sequences) or *E. coli* K12 database (Proteome ID: UP000000625; Downloaded in 10/11/2021 containing 4345 reviewed sequences). Common contaminants were added to the databases. Enzyme specificity was set to trypsin with up to two missed cleavages. For APQ and deep-APQ method, carbamidomethylation (C) (+57.021 Da), peptide N-terminal and lysine dimethyl labeling (+32.05056 Da (32L) and +32.05641 Da (32H)) were set as fixed modification, oxidation (M) (+15.995 Da) was set as variable modifications. For APQ, and OTOT mode of deep-APQ method, the mass tolerances were set to 20 ppm for precursor ions and 20 ppm for fragment ions. For ITOT mode of deep-APQ method, the mass tolerances were set to 20 ppm for precursor ions and 0.5 Da for fragment ions. False discovery rates (FDRs) of 1% for peptide and protein identifications were required. Quantitation analysis was simultaneously performed by home-made quantitative software (deepquant), the intensities of a1 ion clusters of all PSMs were extracted within 20 ppm mass tolerance and used for relative quantitation. Besides, if multiple ions existed in the mass tolerance of a1 ions, the highest intensity was considered as the a1 ion intensity. Peptide and protein ratios were calculated as the median of all the spectra matching the peptide and all quantified peptides matching the protein, respectively.

For LFQ method, all raw files were analyzed together in the MaxQuant environment (version 1.6.5) and employed Andromeda for database search. MS/MS spectra were searched against the human protein database (Proteome ID: UP000005640; Downloaded in 08/10/20 containing 38007 reviewed sequences). Enzyme specificity was set to trypsin with up to two missed cleavages. Oxidation (M) (+15.995 Da) and acetylation (protein N-termini) (+42.011 Da) were set as variable modifications. The mass tolerances were 10 ppm for the precursor ions and 20 ppm for the fragment ions. The time window used in ‘match between runs’ for the transfer of identifications was 1 min. LFQ algorithm was applied for label free protein quantification with at least 2 peptides for a protein. Only unique and razor peptides were used for quantification.

For intensity-based absolute quantification (iBAQ) analysis of *E. coli* in two kinds of growth median, acquired raw files were proposed by Maxquant (version 1.6.5), and searched against *E*.*coli* K12 database (Proteome ID: UP000000625; Downloaded in 10/11/2021 containing 4345 reviewed sequences). The enzyme specificity was set to trypsin with less than two missed cleavages. The search included cysteine carbamidomethylation as a fixed modification and methionine oxidation as a variable modification. Cysteine carbamidomethylation, peptide N-terminal and lysine dimethyl labeling (+32.05056 Da (32L) and +32.05641 Da (32H)) were set as fixed modification. Methionine oxidation was set as variable modifications. The enzyme was trypsin with a maximum of two missed cleavages. The mass tolerances were set to 10 ppm for precursor ions and 0.5 Da for fragment ions. The FDRs for PSM and protein identifications were controlled under 1%. The quantitation results were calculated using iBAQ method, and only unique and razor peptides were used for quantification.

### 2.7. Bioinformatic Analysis

A normalization process was performed so that the log2 median ratio in two channel equaled zero. Differentially expressed proteins were considered as proteins of which the log2 ratios were filtered after student T-test with a p-value cutoff of 0.05 and at least 1.5-fold ratio changes.

## 3. Results and discussion

### 3.1. Principle of deep-APQ method

Considering the information loss of fragment ions in the high mass range and low intensities of fragment ions by APQ method, we developed the deep-APQ method based on segmented tandem mass spectra to improve the quantitation depth. Peptides were first labeled by 32L (^13^CH^2^O, NaBD^3^CN) and 32H (CD^2^O, NaBH^3^CN) with the mass difference of 5.86 mDa between the two channels, thus, the paired precursor ions could be co-fragmented into the same tandem mass spectrum. Then, a two-step MS/MS acquisition was achieved by sequentially acquiring the high mass range for identification and the low mass range to extract the intensities of paired a1 ions for quantitation (Figure 1).

**Figure 1.**
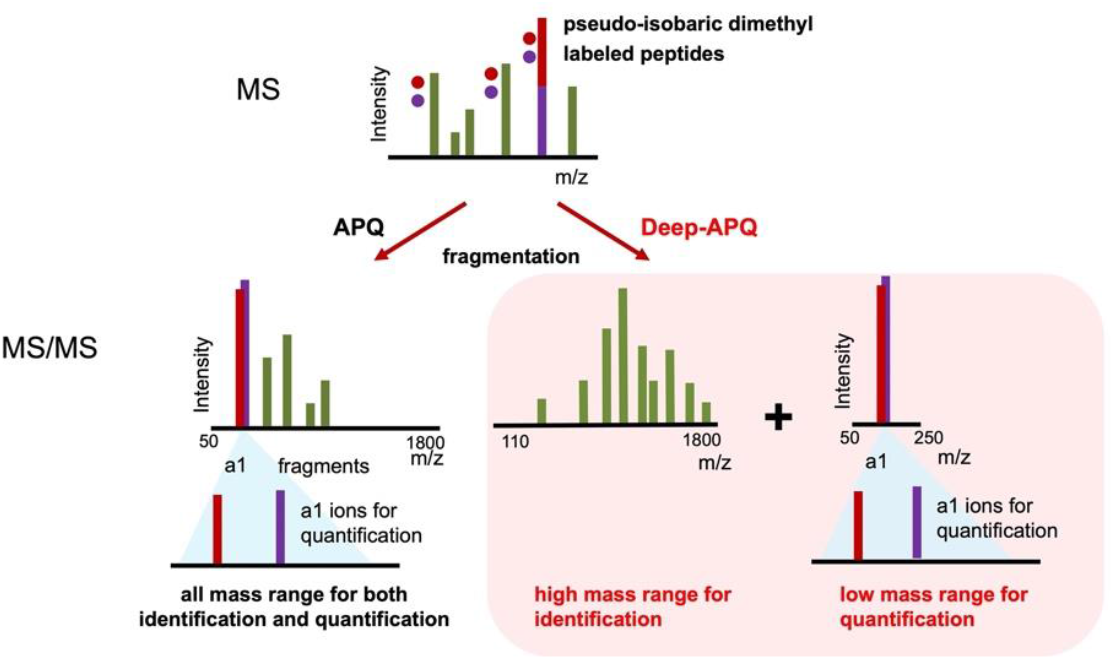
Scheme of deep-APQ method.

The high mass range was set larger than 110 m/z, which was comparable with that of the widely-used label-free method, to guarantee the effective acquisition of large fragment ions and identification of tandem mass spectra. Besides, many high-intensity a1 ions were excluded, reducing the signal masking of some low-abundance fragment ions due to the setting of fixed automatic gain control (AGC). Furthermore, both ion trap (IT) and orbitrap (OT) were compatible as mass analyzers for identification. For the low mass range used for quantification, 50 to 250 m/z was set to include all a1 ions after dimethyl labelling (62 - 191 m/z) and the paired a1 ions were extracted for relative comparison. Therefore, a deep-APQ method was came up to improve proteome coverage by enhancing the S/N ratios of low-abundant fragment ions and the detection of large fragment ions by segmented MS/MS acquisition, which was especially beneficial for the quantitation of long peptides.

### 3.2. Evaluation of quantitation depth

We evaluated the qualitative and quantitative numbers of deep-APQ method using tryptic HeLa protein digests as labeled above. Compared to the APQ method, although the number of acquired MS/MS decreased due to the repeated acquisition of peptides, the identified PSMs increased by 77%, which was because the identification rate of MS/MS increased to 51%. Significant improvements were also shown in the quantitative PSM, peptide and protein numbers by deep-APQ method with the number of peptides increasing by 58%, and the number of proteins increasing by 30% (Supplementary Table 1). Furthermore, by avoiding the unpaired appearance of quantitative information in different MS runs, the deep-APQ method have shown higher quantitation depth than the widely-used label-free quantitation method^17^ (Supplementary Table 2). These results showed that the deep-APQ method would be an effective way to gain a deep quantitation depth for large-scale proteome analysis.

The potential reasons for improved quantitation depth can be attributed to the higher confidence of MS/MS identification and better detection of large fragment ions. As an example, peptide YYGGAEVVDEIELLCQR were detected by both APQ and deep-APQ method. In the tandem mass spectra, more fragment ions were shown in deep-APQ method, especially for the high mass range (Figure 2a, b). The same trend was also shown in other peptides (Supplementary Figure 2). This was because if the first mass was set as low as 50 m/z, fragment ions in the high mass range were poorly transmitted and ion intensities were severely lost due to the limitation of ion transmission from c-trap to orbitrap^18^. Therefore, the q-value, which is a casual way to measure the scoring of spectra, of the deep-APQ method was much lower than that of the APQ method, indicating the higher confidence of MS/MS identification. Furthermore, we calculated the q-values for all PSMs (Figure 2c and Supplementary Figure 3), and the proportion of PSMs with p-value less than 0.01 were 63.26% for deep-APQ method, and 22.43% for APQ method, indicating that more acquired spectrum had higher identification confidence by deep-APQ method. On the other hand, we found that more long peptides were quantified based on the advantage that deep-APQ method can better detect large fragment ions. The percentage of peptides with amino acid number greater than 20 were 7.32% for deep-APQ method and 5.64% for APQ method (Figure 2d). The protein sequence coverage of deep-APQ method (11.22%) was also higher than that of APQ method (9.33%). Therefore, deep-APQ method would be beneficial not only to increase the number of quantified proteins, but also to improve the sequence coverage for each quantified protein.

**Figure 2.**
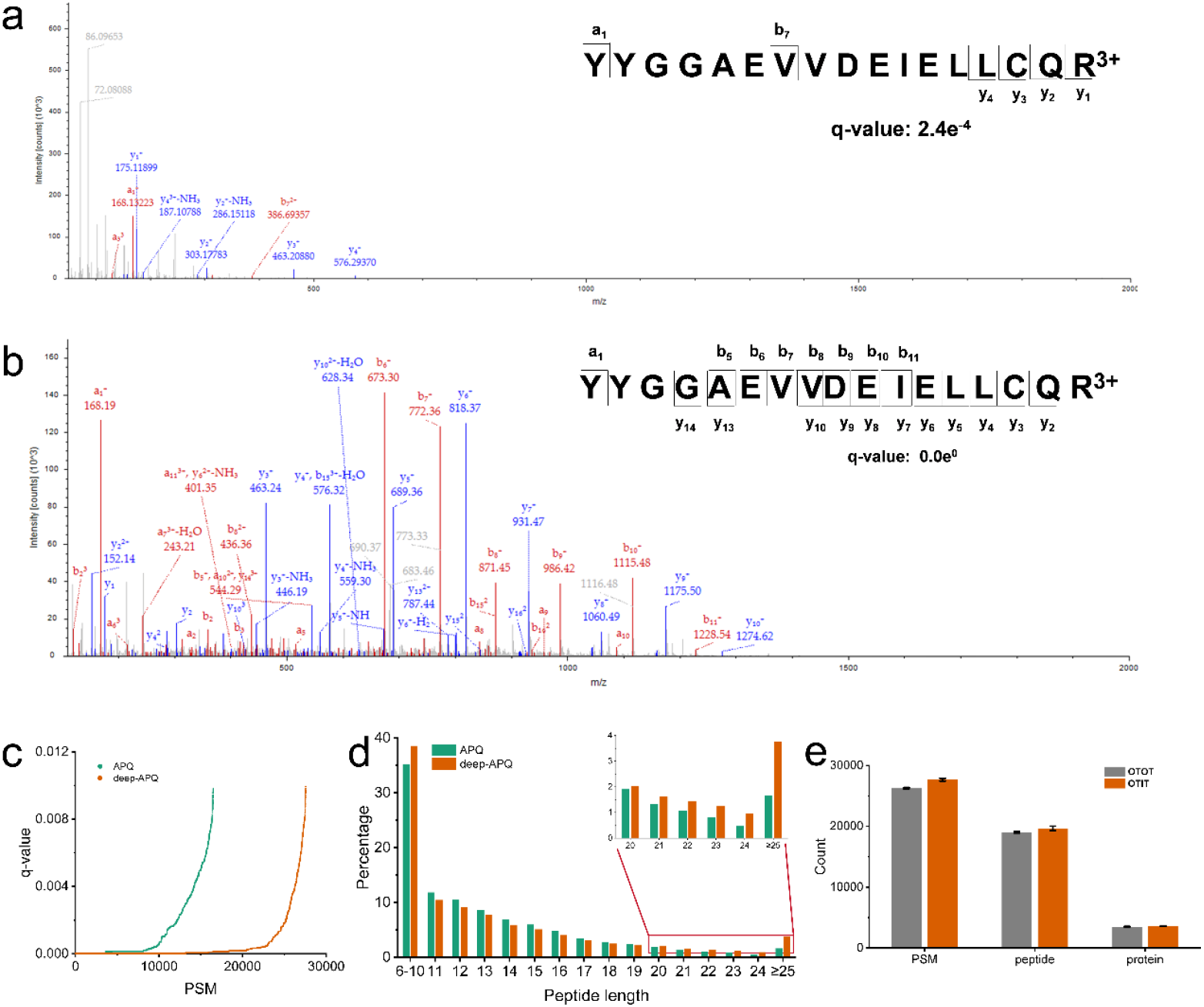
Comparison between APQ and deep-APQ method, and universality of deep-APQ method. An example of the PSM for peptide YYGGAEVVDEIELLCQR acquired in APQ method (a) and deep-APQ method (b). c, Distributions of q-values for all quantitative PSMs acquired by APQ or deep-APQ method. d, Distributions of quantified peptide lengths. e, Quantitation numbers of PSMs, peptides, and proteins acquired using OTOT or ITOT mode in deep-APQ method.

Considering the higher scanning speed of ion trap (IT) than orbitrap (OT), the identification spectra in the high mass range for the above data were acquired using the IT (abbreviated as OTIT). In order to improve the instrument universality of the method, we acquired identification spectra in the high mass range by OT (abbreviated as OTOT) and found they both showed good quantitation depth in terms of the PSM, peptide and protein number, with OTIT a little better than OTOT (Figure 2e). This result demonstrated that the deep-APQ method is versatile, whether applied to the mass spectrometer with two kinds of mass analyzers (such as Orbitrap Fusion Lumos), or the single mass analyzer of orbitrap (such as Q Exactive Orbitrap or Orbitrap Exploris 480).

### 3.3. Quantification accuracy, precision and dynamic range of deep-APQ

Since the APQ method^14^ has shown significantly higher accuracy and precision than reporter ion-based methods, we evaluated whether the segmented acquisition still exhibited the advantages. The 32L and 32H labeled tryptic HeLa protein digests were mixed at the ratios of 1:1, 1:2, 1:5, 1:10 and 1:20, and all experiments were repeated three times. The average of the measured median values were 1:1.3, 1:2.3, 1:4.9, 1:9.6 and 1:20.8 respectively, with the average relative error of 7.4% (Figure 3a). Besides, deep-APQ showed good precision with the average CV value of 6.62% and no significant difference with different mixing ratios (Figure 3b), maintaining the excellent quantitative performance of the APQ method. As the mixing ratio increasing, the number of quantitation proteins and quantitation rate were constant (Figure 3c and Supplementary Table 3). Therefore, the deep-APQ method exhibited excellent accuracy, precision and high quantitation depth within the 20-fold dynamic range, heralding great potential for broad application in quantitative proteome studies.

**Figure 3.**
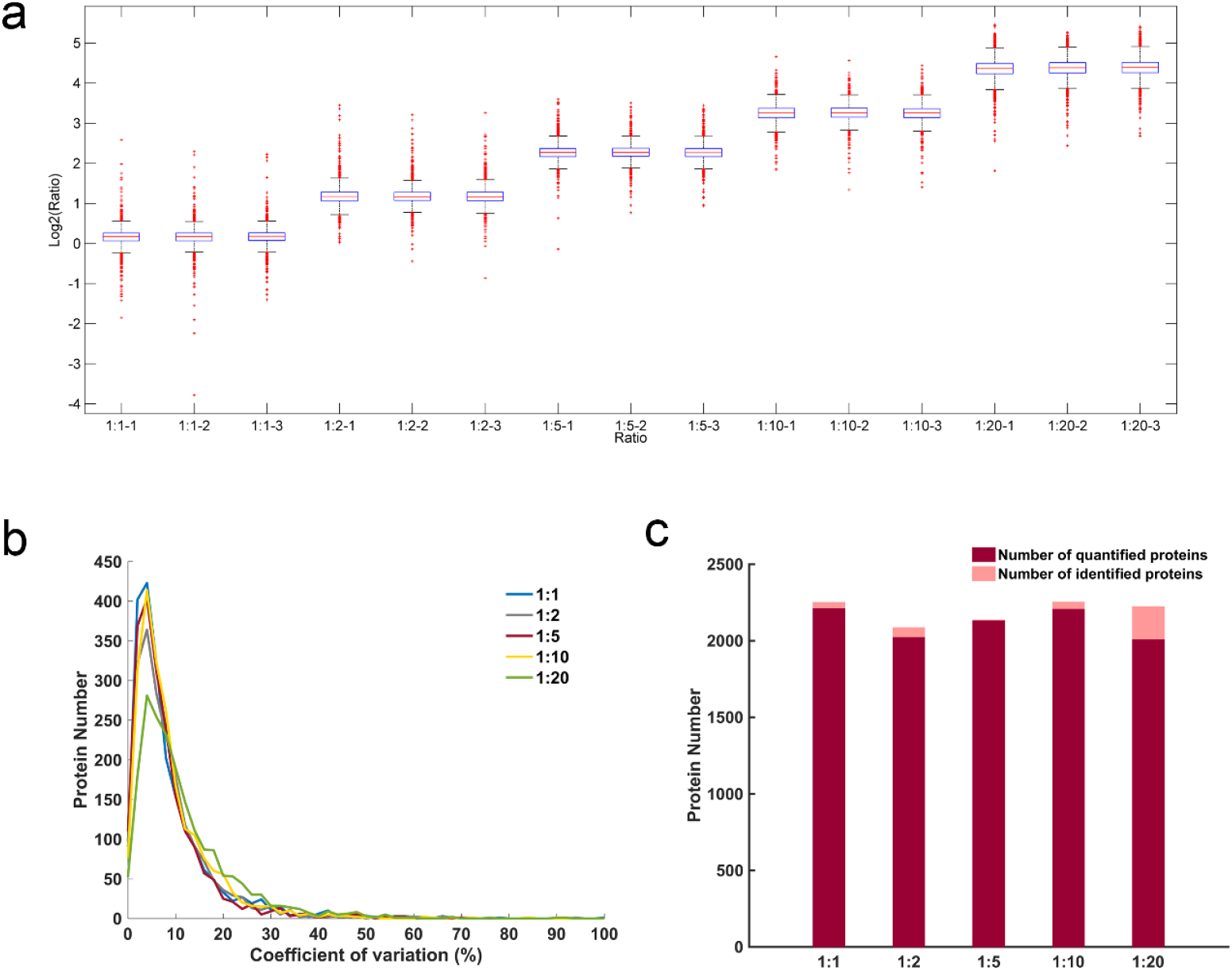
Quantitative evaluation of the deep-APQ method. a, Quantitation results of human proteins mixed at the ratio of 1:1, 1:2, 1:5, 1:10, and 1:20. Box plots showed the median (centred red line), first and third quartiles (lower and upper box limits, respectively), 1.5 times the interquartile range (whiskers) and outliers (cross). b, Coefficients of variation for quantitation ratios of 1:1, 1:2, 1:5, 1:10, and 1:20 mixed human proteins among triplicate LC-MS/MS runs. c, Qualitative and quantitative numbers of human proteins at different mixing ratios.

### 3.4. In-depth quantification of *E. coli* proteomes cultivating in glucose and acetate

How microorganisms organize biological processes to ensure growth and maintenance under different conditions is a basic question in synthetic biology, with important implications for engineering microbial cell factories to produce chemicals and materials precisely and efficiently. However, due to the wide range of protein expression abundances and complexity of pathways, in-depth proteomic mapping in response to varying carbon sources is still urgently needed to elucidate the underlying molecular mechanisms for cell survival. Herein, we applied the deep-APQ method to study the dynamic *E. coli* proteomes under the carbon sources of glucose and acetate. Glucose is a traditional carbon source which is easily utilized by *E. coli*. On the other hand, acetate is the fermentation product of *E. coli* with toxicity and poor energy content^19, 20^, but has been used as a cheap and abundant feedstock to produce recombinant protein^21^, biofuel^22^, or biochemical^23^. We harvested cells at mid-exponential growth phase separately with three biological replicates (Supplementary Figure 1).

To achieve deep proteome analysis, we integrated a workflow combining deep-APQ with highly efficient protein extraction^24^ and three-dimensional (3D) separation (Figure 4a). The 3D separation including high-pH (HpH) and low-pH reversed-phase separation and high-field asymmetric waveform ion mobility spectrometry (FAIMS) were used to reduce sample complexity and increase the protein dynamic range. Especially, as an important parameter, we optimized the voltages of FAIMS and used the combination of 40V, 60V and 80V to get the largest quantified number of proteins and smallest protein overlap (Supplementary Figure 4). In total, 3447 *E. coli* proteins, with average protein sequence coverage of 40%, were quantified, representing 82% of gene-coding proteins^25^, showing a deeper proteome landscape than the reported quantitative proteome works ^26, 27^. The quantitative data had high reproducibility with the median CV of 17%. The absolute abundance of quantified proteins spaned 6 orders of magnitude referring to the method of intensity-based absolute quantification (iBAQ), covering some low-abundance proteins enriched in biological processes of bacterial-type flagellum-dependent cell motility, chemotaxis, and transmembrane transport (Figure 4b). Besides, the quantified proteins were localized in various cell compositions, such as ribosome, cell outer membrane, bacterial-type flagellum (Figure 4c).

**Figure 4.**
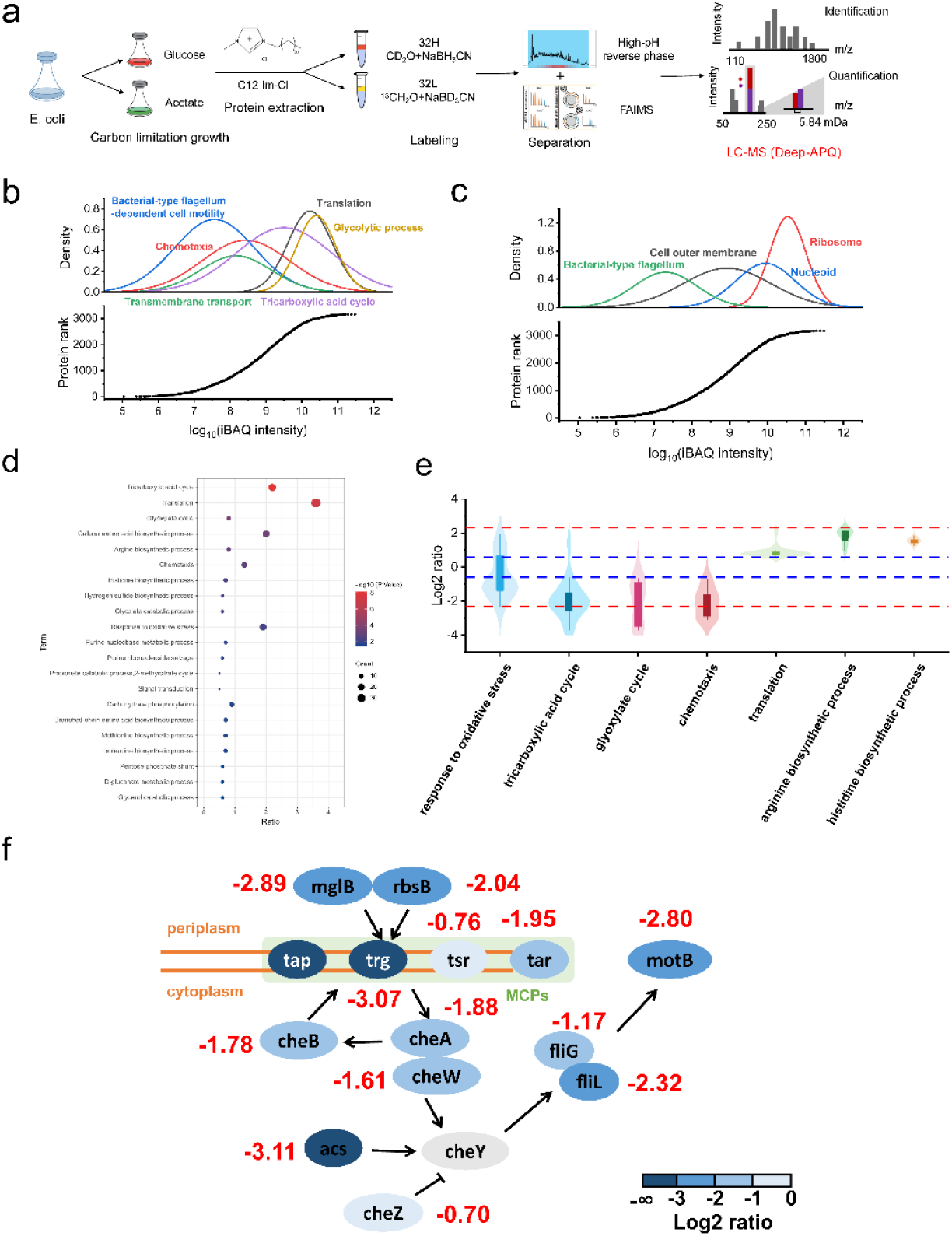
In-depth quantitation in *E*.*coli* system. a, An in-depth workflow of the impact of carbon sources on *E. coli* proteomics. b, IBAQ intensity distribution of different biological processes. c, IBAQ intensity distribution of different cell composition. d, Main biological process of GO analysis. e, Violin diagram of ratio distribution of different biological processes. f, Network of quantified proteins associated with the chemotaxis process.

Considering the significant difference of growth rates under the two growth conditions (Supplementary Figure 1), we further study the changes of its metabolic model. A total of 1074 proteins were significantly regulated with 487 up-regulated and 587 down-regulated in glucose media comparing to acetate media. They were mainly involved in the tricarboxylic acid cycle, translation, glyoxylic acid cycle and cellular amino acid biosynthesis (Figure 4d). In particular, many proteins enriched in different processes showed a collective upward or downward trend basically (Figure 4e). For example, AceA, aceB, ppsA and pckA played an important role in glyoxylate process as gluconeogenic enzymes^16^, and showed an up-grade trend in acetate media. While in glucose media, the enzymes in cellular amino biosynthetic process including hisC, leuC, and trpGD, were highly expressed, indicating that the *E. coli* cells were in a stable state of protein synthesis and production, rather than trying to supply energy as in acetate medium.

Due to the deep coverage of our dataset, we not only observed the adjustment of metabolism under the two carbon sources, but also the changes to induce stress protection and tactic behavior. In particular, extensive down-regulations of proteins related to chemotaxis with low abundances was found in glucose media (Figure 4f). As one of the most complex and costly bacterial behaviors, chemotaxis was an important mechanism to enhance adaptability in poor nutritional conditions at the cost of reducing the investment in the growth trade-off^28^.In our data, mglB, rbsB, tap, trg, tsR, taR, cheA, cheB, cheW, liM, fliG, fliL, and motB showed higher expressions in acetate growth media, showing a strong chemotaxis performance in the poor nutritional condition. Besides, another chemotactic enzyme acs, an essential intermediate at the junction of anabolic and catabolic pathways, enables the cell to use acetate during aerobic growth to generate energy via the TCA cycle, and is the response regulator involved in flagellar movement and chemotaxis, also show the high expression in acetate growth media. Our in-depth quantitative *E. coli* landscape showed the detailed protein dynamics of chemotaxis to understand the stress behavior, hoping to help the reduction of the effects of acetate toxicity for synthetic biology and metabolic engineering.

## 4. Conclusions

In summary, we developed an in-depth proteome quantitation method, deep-APQ, based on segmented tandem mass spectra acquisition of a1 ion-based quantitation. Benefit from a greater number of fragment ions with high mass and efficient removal of many high-abundance a1 ions, the depth of quantitative proteins was higher than the traditional APQ method and the widely-used label-free quantitation method. Besides, it still showed high quantitation accuracy and precision within 20-fold dynamic range. We integrated a workflow to achieve an in-depth quantitation proteome analysis of *E. coli* responses to different carbon sources and observed the significant up-regulations of proteins related to the low-abundance chemotaxis process in the poor nutritional acetate condition. Deep-APQ method shows great potentials to draw global landscapes of the dynamic proteomes for biomedicine, neuroscience or metabolic engineering studies with high depth and accuracy.

## Supporting information

Considering the significant difference of growth rates under the two growth conditions

## Acknowledgements

The authors are grateful for the financial support from National Key R&D Program of China (2017YFA0505003), National Natural Science Foundation (22104138), Project funded by China Postdoctoral Science Foundation (2020M670804), United Innovation Program from DICP and QIBEBT, CAS (DICP&QIBEBT UN201803), Talent innovation support program of Dalian (2019CT07).

## References

1. Angel, T. E.; Aryal, U. K.; Hengel, S. M.; Baker, E. S.; Kelly, R. T.; Robinson, E. W.; Smith, R. D. Mass spectrometry-based proteomics: existing capabilities and future directions. Chemical Society Reviews 2012, 41,10, 3912.

2. Post, H.; Penning, R.; Fitzpatrick, M. A.; Garrigues, L. B.; Wu, W.; MacGillavry, H. D.; Hoogenraad, C. C.; Heck, A. J. R.; Altelaar, A. F. M. Robust, Sensitive, and Automated Phosphopeptide Enrichment Optimized for Low Sample Amounts Applied to Primary Hippocampal Neurons. Journal of Proteome Research 2016, 16,2, 728–737.

3. Mehta, S.; Easterly, C. W.; Sajulga, R.; Millikin, R. J.; Argentini, A.; Eguinoa, I.; Martens, L.; Shortreed, M. R.; Smith, L. M.; McGowan, T.; Kumar, P.; Johnson, J. E.; Griffin, T. J.; Jagtap, P. D. Precursor Intensity-Based Label-Free Quantification Software Tools for Proteomic and Multi-Omic Analysis within the Galaxy Platform. Proteomes 2020, 8,3.

4. Zhang, F.; Ge, W.; Ruan, G.; Cai, X.; Guo, T. Data-Independent Acquisition Mass Spectrometry-Based Proteomics and Software Tools: A Glimpse in 2020. Proteomics 2020, 20,17-18, e1900276.

5. Shah, A. D.; Goode, R. J. A.; Huang, C.; Powell, D. R.; Schittenhelm, R. B. LFQ-Analyst: An Easy-To-Use Interactive Web Platform To Analyze and Visualize Label-Free Proteomics Data Preprocessed with MaxQuant. J Proteome Res 2020, 19,1, 204–211.

6. Borràs, E.; Sabidó, E. DIA+: A Data -Independent Acquisition Method Combining Multiple Precursor Charges to Improve Peptide Signal. Analytical Chemistry 2018, 90,21, 12339–12341.

7. Li, J.; Van Vranken, J. G.; Pontano Vaites, L.; Schweppe, D. K.; Huttlin, E. L.; Etienne, C.; Nandhikonda, P.; Viner, R.; Robitaille, A. M.; Thompson, A. H.; Kuhn, K.; Pike, I.; Bomgarden, R. D.; Rogers, J. C.; Gygi, S. P.; Paulo, J. A. TMTpro reagents: a set of isobaric labeling mass tags enables simultaneous proteome-wide measurements across 16 samples. Nature methods 2020, 17,4, 399–404.

8. Ross, P. L.; Huang, Y. N.; Marchese, J. N.; Williamson, B.; Parker, K.; Hattan, S.; Khainovski, N.; Pillai, S.; Dey, S.; Daniels, S.; Purkayastha, S.; Juhasz, P.; Martin, S.; Bartlet-Jones, M.; He, F.; Jacobson, A.; Pappin, D. J. Multiplexed protein quantitation in Saccharomyces cerevisiae using amine-reactive isobaric tagging reagents. Mol. Cell. Proteomics 2004, 3,12, 1154–69.

9. Hogrebe, A.; von Stechow, L.; Bekker-Jensen, D. B.; Weinert, B. T.; Kelstrup, C. D.; Olsen, J. V. Benchmarking common quantification strategies for large-scale phosphoproteomics. Nature communications 2018, 9,1, 1045.

10. Liu, J.; Zhou, Y.; Shan, Y.; Zhao, B.; Hu, Y.; Sui, Z.; Liang, Z.; Zhang, L.; Zhang, Y. A Multiplex Fragment-Ion-Based Method for Accurate Proteome Quantification. Anal. Chem. 2019, 91,6, 3921–3928.

11. Nie, A. Y.; Zhang, L.; Yan, G. Q.; Yao, J.; Zhang, Y.; Lu, H. J.; Yang, P. Y.; He, F. C. In vivo termini amino acid labeling for quantitative proteomics. Anal. Chem. 2011, 83,15, 6026–33.

12. Tian, X.; de Vries, M. P.; Visscher, S. W. J.; Permentier, H. P.; Bischoff, R. Selective Maleylation-Directed Isobaric Peptide Termini Labeling for Accurate Proteome Quantification. Anal. Chem. 2020, 92,11, 7836–7844.

13. Waldbauer, J.; Zhang, L.; Rizzo, A.; Muratore, D. diDO-IPTL: A Peptide-Labeling Strategy for Precision Quantitative Proteomics. Anal. Chem. 2017, 89,21, 11498–11504.

14. Liu, J.; Zhou, Y.; Hou, X.; Liu, C.; Zhao, B.; Shan, Y.; Sui, Z.; Liang, Z.; Zhang, L.; Zhang, Y. A1 Ions: Peptide-Specific and Intensity-Enhanced Fragment Ions for Accurate and Multiplexed Proteome Quantitation. Analytical Chemistry 2022. 94, 7637–7646.

15. Csonka, L. N., Ikeda, T. P., Fletcher, S. A. & Kustu, S. The accumulation of glutamate is necessary for optimal growth of Salmonella typhimurium in media of high osmolality but not induction of the proU operon. J. Bacteriol. [J] 1994, 176, 6324–6333

16. Basan, M.; Honda, T.; Christodoulou, D.; Hörl, M.; Chang, Y. -F.; Leoncini, E.; Mukherjee, A.; Okano, H.; Taylor, B. R.; Silverman, J. M.; Sanchez, C.; Williamson, J. R.; Paulsson, J.; Hwa, T.; Sauer, U. A universal trade-off between growth and lag in fluctuating environments. Nature 2020, 584,7821, 470–474.

17. Cox, J.; Hein, M. Y.; Luber, C. A.; Paron, I.; Nagaraj, N.; Mann, M. Accurate proteome-wide label-free quantification by delayed normalization and maximal peptide ratio extraction, termed MaxLFQ. Mol. Cell. Proteomics 2014, 13,9, 2513–26.

18. Vasicek, L. A.; Ledvina, A. R.; Shaw, J.; Griep-Raming, J.; Westphall, M. S.; Coon, J. J.; Brodbelt, J. S. Implementing Photodissociation in an Orbitrap Mass Spectrometer. Journal of the American Society for Mass Spectrometry 2011, 22,6, 1105–1108.

19. Oh, M.-K.; Rohlin, L.; Kao, K. C.; Liao, J. C. Global Expression Profiling of Acetate-grown Escherichia coli. Journal of Biological Chemistry 2002, 277,15, 13175–13183.

20. Polen, T.; Rittmann, D.; Wendisch, V. F.; Sahm, H. DNA Microarray Analyses of the Long-Term Adaptive Response of Escherichia coli to Acetate and Propionate. Applied and Environmental Microbiology 2003, 69,3, 1759–1774.

21. Leone, S.; Sannino, F.; Tutino, M. L.; Parrilli, E.; Picone, D. Acetate: friend or foe? Efficient production of a sweet protein in Escherichia coli BL21 using acetate as a carbon source. Microbial Cell Factories 2015, 14,1.

22. Song, H.-S.; Seo, H.-M.; Jeon, J.-M.; Moon, Y.-M.; Hong, J. W.; Hong, Y. G.; Bhatia, S. K.; Ahn, J.; Lee, H.; Kim, W.; Park, Y.-C.; Choi, K.-Y.; Kim, Y.-G.; Yang, Y.-H. Enhanced isobutanol production from acetate by combinatorial overexpression of acetyl-CoA synthetase and anaplerotic enzymes in engineered Escherichia coli. Biotechnology and Bioengineering 2018, 115,8, 1971–1978.

23. Huang, B.; Yang, H.; Fang, G.; Zhang, X.; Wu, H.; Li, Z.; Ye, Q. Central pathway engineering for enhanced succinate biosynthesis from acetate inEscherichia coli. Biotechnology and Bioengineering 2018, 115,4, 943–954.

24. Fang, F.; Zhao, Q.; Chu, H.; Liu, M.; Zhao, B.; Liang, Z.; Zhang, L.; Li, G.; Wang, L.; Qin, J.; Zhang, Y. Molecular Dynamics Simulation-assisted Ionic Liquid Screening for Deep Coverage Proteome Analysis. Molecular & Cellular Proteomics 2020, 19,10, 1724–1737.

25. Bartholomäus, A.; Fedyunin, I.; Feist, P.; Sin, C.; Zhang, G.; Valleriani, A.; Ignatova, Z. Bacteria differently regulate mRNA abundance to specifically respond to various stresses. Philosophical Transactions of the Royal Society A: Mathematical, Physical and Engineering Sciences 2016, 374,2063, 20150069.

26. Navarrete-Perea, J.; Gygi, S. P.; Paulo, J. A. Growth media selection alters the proteome profiles of three model microorganisms. Journal of proteomics 2021, 231, 104006.

27. Mori, M.; Zhang, Z.; Banaei-Esfahani, A.; Lalanne, J. B.; Okano, H.; Collins, B. C.; Schmidt, A.; Schubert, O. T.; Lee, D. S.; Li, G. W.; Aebersold, R.; Hwa, T.; Ludwig, C. From coarse to fine: the absolute Escherichia coli proteome under diverse growth conditions. Molecular systems biology 2021, 17,5.

28. Ni, B.; Colin, R.; Link, H.; Endres, R. G.; Sourjik, V. Growth-rate dependent resource investment in bacterial motile behavior quantitatively follows potential benefit of chemotaxis. Proceedings of the National Academy of Sciences 2019, 117,1, 595–601.

