## Supplementary material for "Segmented MS/MS acquisition of a1 ion-based strategy for in-depth proteome quantitation": Considering the significant difference of growth rates under the two growth conditions

**Supplementary Table 1 | Quantitative results of deep-APQ and APQ method**

| | MS/MS | Identification PSMs | Quantitative PSMs | Quantitative peptides | Quantitative proteins |
|---|---|---|---|---|---|
| **deep-APQ** | 58575 | 29589 | 27661 | 19665 | 3586 |
| **APQ** | 68985 | 16378 | 16236 | 12314 | 2690 |

**Supplementary Table 2 | Quantitation protein numbers of deep-APQ method and label-free method.**

| | Replicate 1 | Replicate 2 | Replicate 3 |
|---|---|---|---|
| **deep-APQ** | 2215 | 2226 | 2198 |
| **Label free** | 1491 | 1641 | 1661 |

**Supplementary Table 3 | Quantitative proteins in different ratios of three technical replicates.**

| Sample | 1:1 | 1:2 | 1:5 | 1:10 | 1:20 |
|---|---|---|---|---|---|
| **r1** | 2215 | 1971 | 2119 | 2216 | 2015 |
| **r2** | 2226 | 2026 | 2132 | 2195 | 1983 |
| **r3** | 2198 | 2081 | 2152 | 2219 | 2033 |

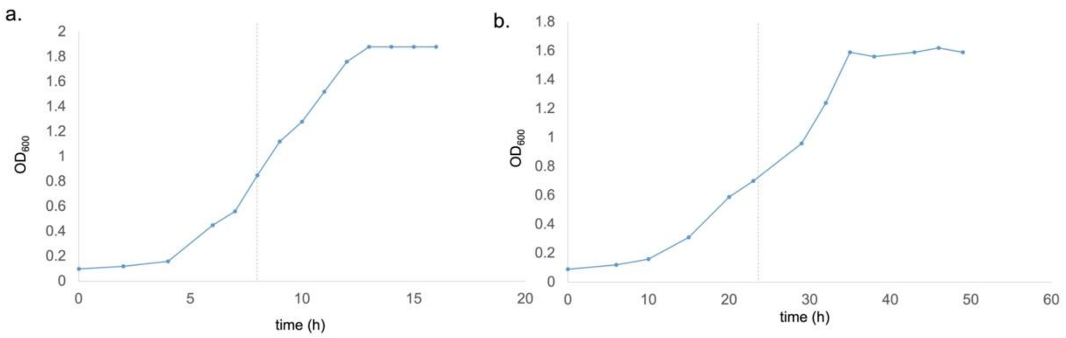


**Supplementary Figure 1 | Growth curve of *E. coli* in glucose (a) and acetate (b) as the only carbon source in the minimum growth media.**

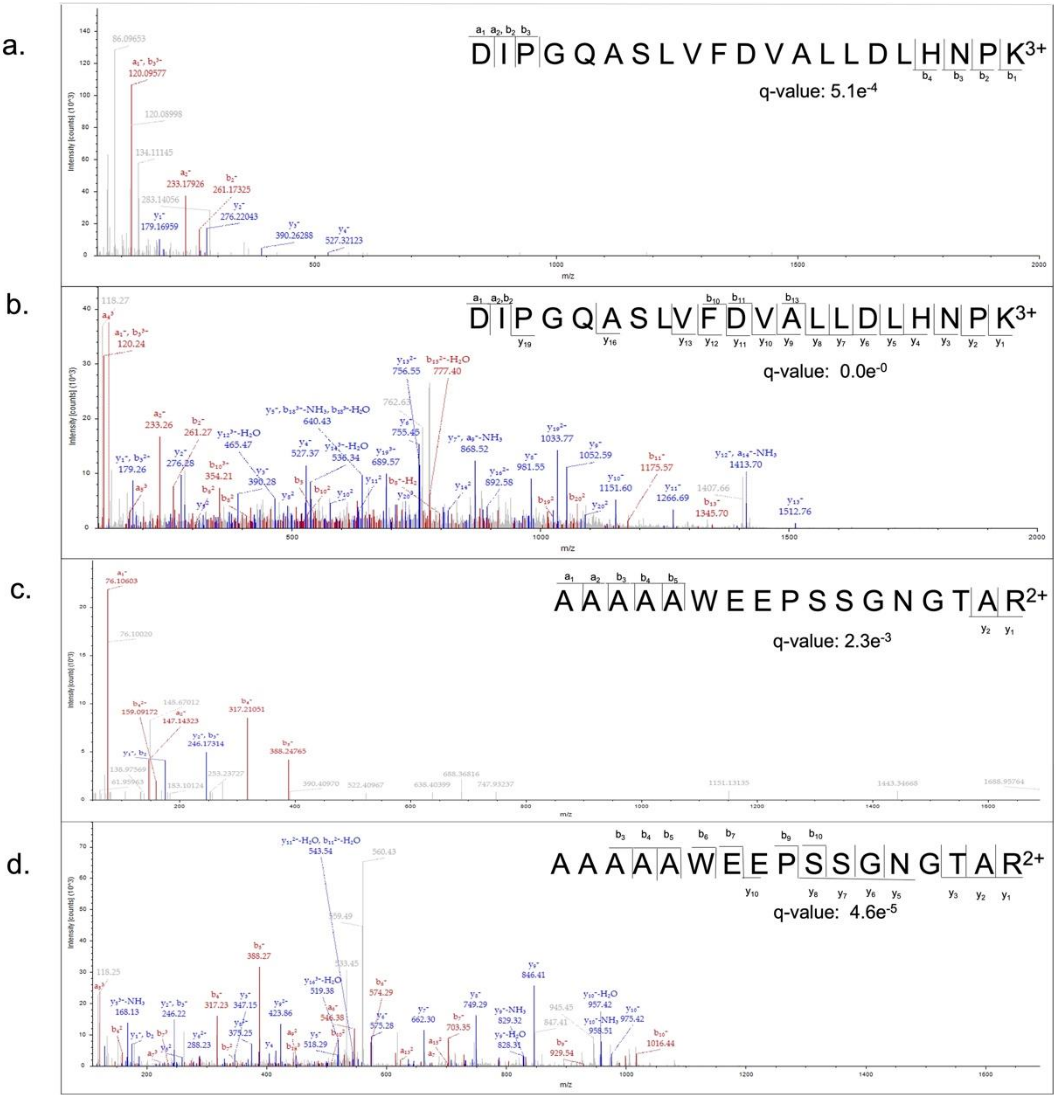


**Supplementary Figure 2 | Other randoms examples of PSM inferred to peptides acquired by APQ or deep-APQ. Peptide DIPGQASLVFDVALLDLHPK acquired by APQ (a) and deep-APQ (b). Peptide AAAAAWEEPSSGNGTAR acquired by APQ (c) and deep-APQ (d).**

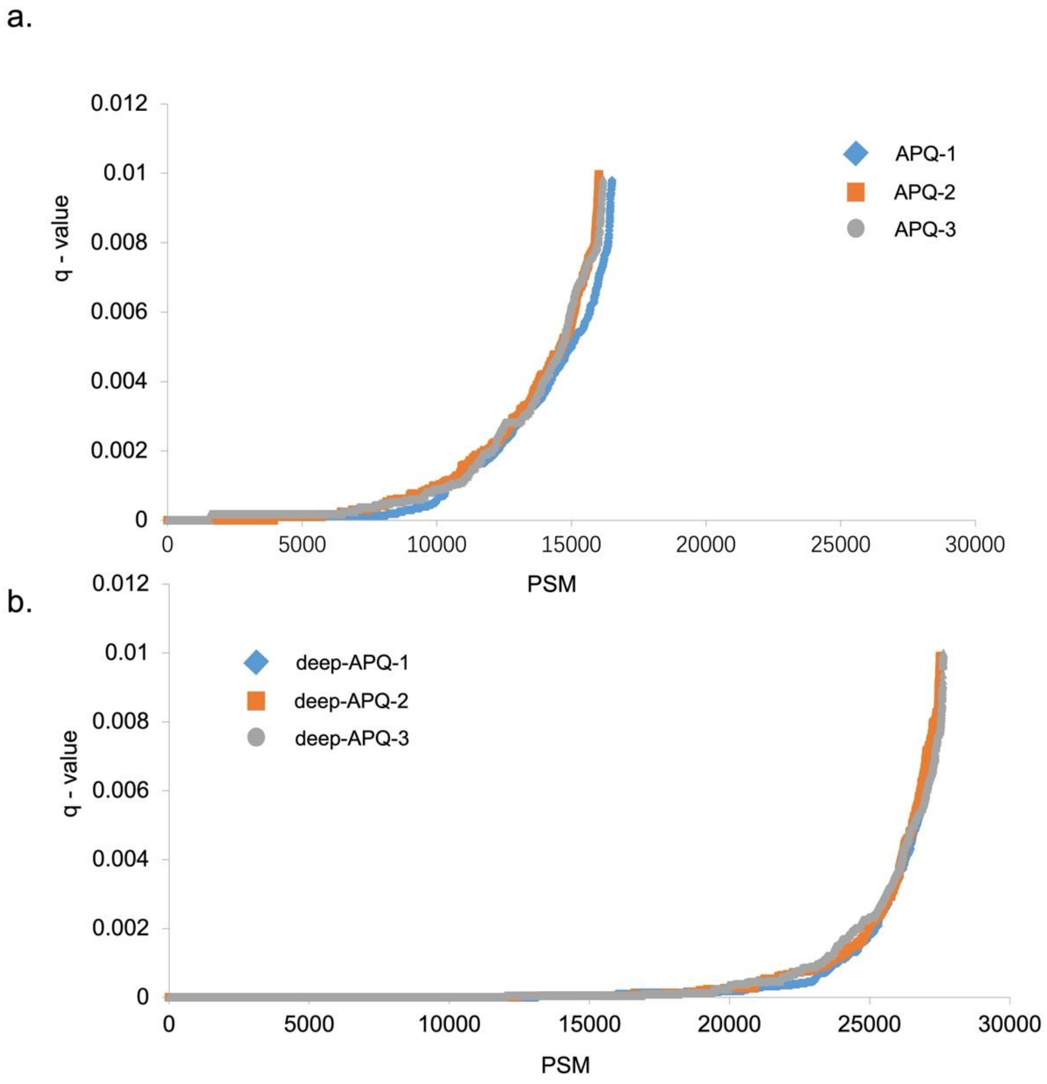


**Supplementary Figure 3 | Q-value distribution of three technical replicates in APQ (a) and deep-APQ (b) acquisition modes.**

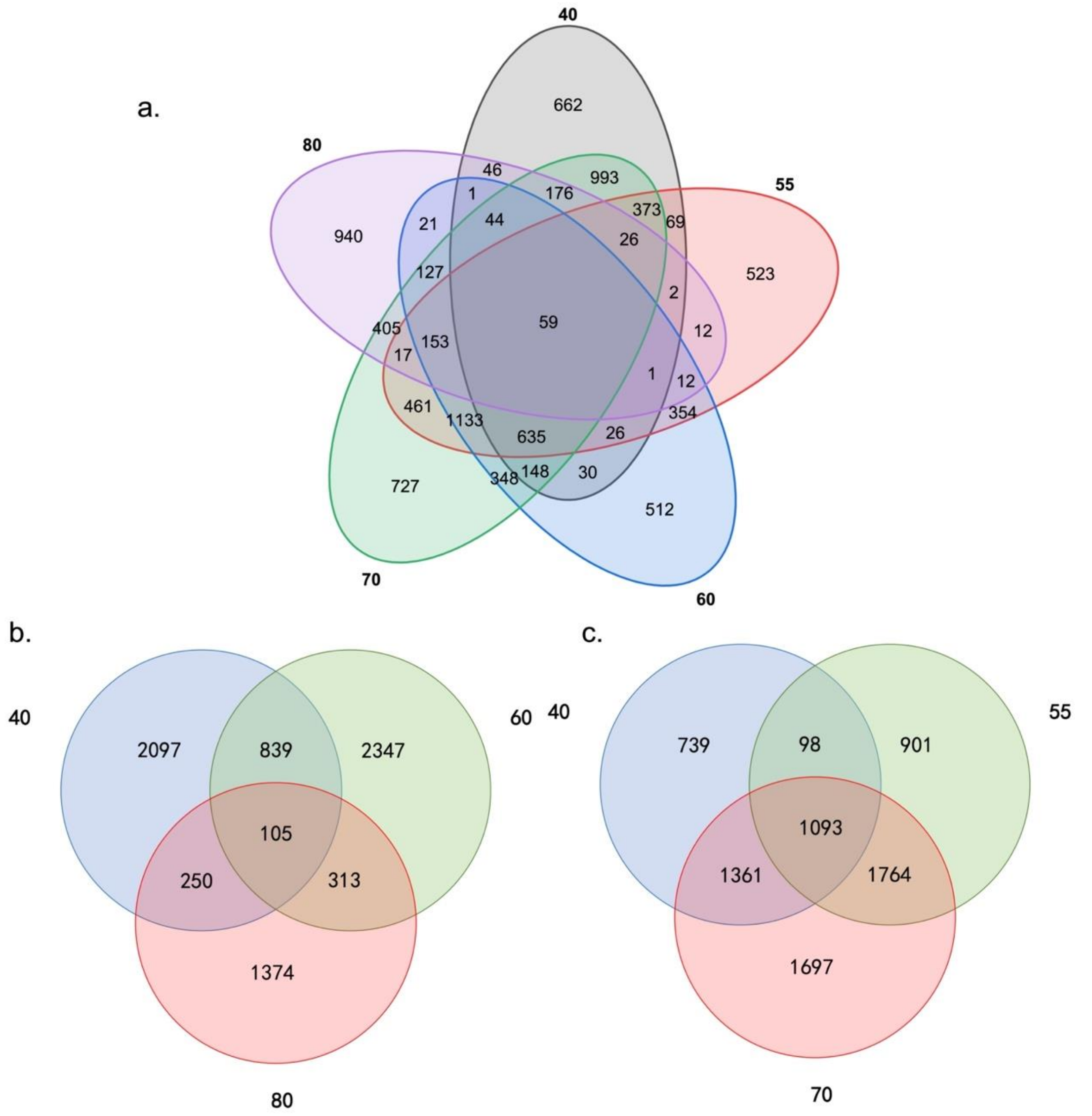


**Supplementary Figure 4 | E.coli proteins identified in different FAIMS voltages (a), the combination of 40, 60, and 80V (b), and the combination of 40, 55, and 70V (c).**
